# Shape Analysis of the Human Association Pathways

**DOI:** 10.1101/2020.04.19.049544

**Authors:** Fang-Cheng Yeh

## Abstract

Shape analysis has been widely used in digital image processing and computer vision, but they have not been utilized to compare the structural characteristics of the human association pathways. Here we used shape analysis to derive length, area, volume, and shape metrics from diffusion MRI tractography and utilized them to study the morphology of human association pathways. The reliability analysis showed that shape descriptors achieved moderate to good test-retest reliability. Further analysis on association pathways showed left dominance in the arcuate fasciculus, cingulum, uncinate fasciculus, frontal aslant tract, and right dominance in the inferior fronto-occipital fasciculus and inferior longitudinal fasciculus. The superior longitudinal fasciculus has a mixed lateralization profile with different metrics showing either left or right dominance. The analysis of between-subject variations shows that the overall layout of the association pathways does not variate a lot across subjects, as shown by low between-subject variation in length, span, diameter, and radius. In contrast, the area of the pathway innervation region has a considerable between-subject variation. A follow-up analysis is warranted to thoroughly investigate the nature of population variations and their structure-function correlation.

## Introduction

Deciphering the structural layout of the human brain has been a challenging goal to understand how structure defines the brain function (DeFelipe, 2010). The first connectome study identified structural connection using diffusion MRI fiber tracking (Sporns et al., 2005) and formulated brain connections as a graph to reveal the network topology (Bullmore and Sporns, 2009). Further studies have correlated structural connectivity with brain function in the healthy population or disease population (Fornito et al., 2015). The network analysis tackled the structure-function correlation from a panoramic view, but the shape characteristics and morphology of the connecting bundles were mostly ignored, particularly the association pathways in the human brain that control most of the cognitive functions. On the other hand, existing tractography studies only used basic shape features such as volume or size and discarded rich morphology information in the fiber pathways (Abhinav et al., 2014; Huang et al., 2005; Lopez et al., 2013; Wolff et al., 2015). More advanced shape analysis focused on specific applications, (Bastin et al., 2008; Corouge et al., 2004; Glozman et al., 2018; Kitchell et al., 2018) and did not provide a comprehensive shape analysis to exploit the all morphology information. There is yet a comprehensive study utilizing shape analysis to investigate the structural characteristics of the human association pathways.

Here we aim to bridge this information gap by applying a comprehensive shape analysis, including length, area, volume, and shape metrics, to investigate the shape characteristics of the human association pathways. Shape analysis has been widely used in computer vision in a variety of applications to achieve imaging understanding of an object (Costa and Cesar Jr, 2000; Russ, 2002). The analysis provides the “shape descriptor”—a quantitative measurement that describes one part of the shape characteristics as length, area, and volume. Leveraging shape analysis to investigate tractography, however, faces two technical challenges. First, the existing shape analysis is designed for 2D pixel-based or 3D voxel-based images, whereas tractography is a set of coordinate sequences plotting the simulated routes of brain connections. The definition of shape descriptors, such as length, area, and volume metrics, requires a substantial revision to fit into the tractography context. Second, the reproducibility of tractography has long been an ongoing issue (Rheault et al., 2020). Without a reliable and reproducible tractography input, the result of shape analysis will be meaningless due to “garbage in, garbage out.”

In this study, we utilized “augmented fiber tracking” to tackle the reproducibility issue and to provide automatic track recognition. Augmented fiber tracking includes three strategies—parameter saturation, atlas-based track recognition/filtering, and topology-informed pruning. “Parameter saturation” was done by tracking millions of tracks using a random combination of anisotropy threshold, step size, and angular threshold to saturate the parameter space. This approach explored millions of parameter combinations to maximize the mapping of fiber pathways. The generated tracks were further recognized and filtered using an expert-vetted tractography atlas (Yeh et al., 2018). This automatic recognition method isolated target pathways and simultaneously excluded irrelevant or false connections that substantially deviated from the known trajectories. After recognition, we further applied topology-informed pruning (Yeh et al., 2019) to eliminate possible false connections. TIP used track density at each voxel to eliminate noisy tracks that failed to form a bundle. We integrated these three strategies to map 14 association pathways on a test-retest dataset from the human connectome projects (n=44).

Then we introduced the shape descriptors for the tractography. Figure 1 illustrates the calculation using the left arcuate fasciculus as an example. Figure 1a shows the quantification of the length metrics, including length, span, diameters of the bundle, and radius of the innervation regions. Figure 1b shows the area metrics, including the area of the entire track surface and area of the two end surfaces. Figure 1c shows the volume metrics, including total track volume and trunk volume. Based on these metrics, we further derived “shape metrics,” which are unit free indices, including curl, elongation, and irregularity, to describe the shape characteristics of the association pathways. We examined the reliability of these metrics using the intra-class correlations (ICC). This reliability results allowed us to identify findings with poor reproducibility and to ensure the robustness of the results. Then we derived the distributions of shape descriptors to reveal their left-right asymmetry and between-subject variations to study the morphology of the association pathways.

**Figure 1.**
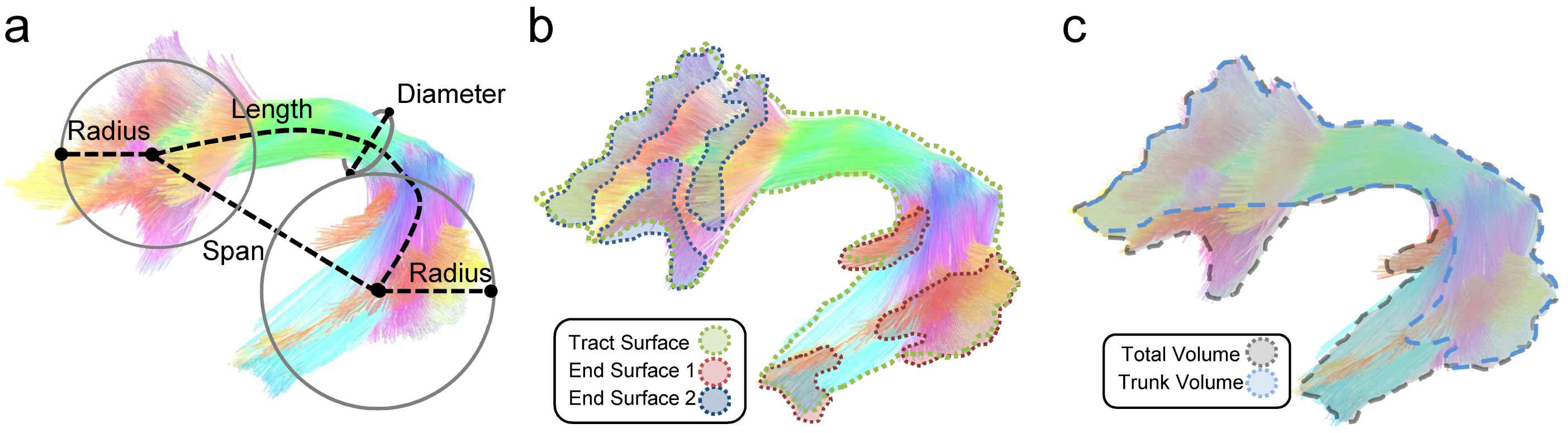
Shape analysis of a bundle. (a) The length metrics include length, span, diameter, and radius of the innervation region. The length measures the length of the bundle trajectory, whereas the span measures the absolute distance between two ends of the bundle. The diameter estimates the average bundle diameter. The radius uses a circular model to estimate the coverage of the innervation regions. (b) The area metrics include total track surface area and area of the two end surfaces. Each fiber bundle has two end surfaces, and their area will be quantified separately. (c) The volume metrics include total volume and trunk volume.

## Material and Methods

### MRI acquisitions

The test-retest diffusion MRI data were acquired from the Human Connectome Project database (WashU consortium)(Glasser et al., 2016). A total of 44 subjects had repeat diffusion MRI scans. 24 of them were female, and 20 of them were male. The age range was 22-to 35-year-old, and the average age was 30.3. One subject was left-handed. The data were acquired using a multishell diffusion scheme with three b-values at 1000, 2000, and 3000 s/mm^2^. Each shell had 90 sampling directions. The spatial resolution was 1.25 mm isotropic. The acquisition parameters are detailed in the consortium paper (Glasser et al., 2016).

### Diffusion MRI fiber tracking

The diffusion data were first rotated and interpolated to the ICBM2009 T1W template at 1mm. Here the rotation used a rigid-body transformation without a nonlinear deformation so that the shape features were preserved. The b-table was also rotated accordingly. The purpose of this spatial transformation was to facilitate a direct comparison of the tractography between the repeat scans. The rotated data were then reconstructed using generalized q-sampling imaging (Yeh et al., 2010) with a diffusion sampling length ratio of 1.7. The b-table was checked by an automatic quality control routine to ensure its accuracy (Schilling et al., 2019).

**Table 1:**
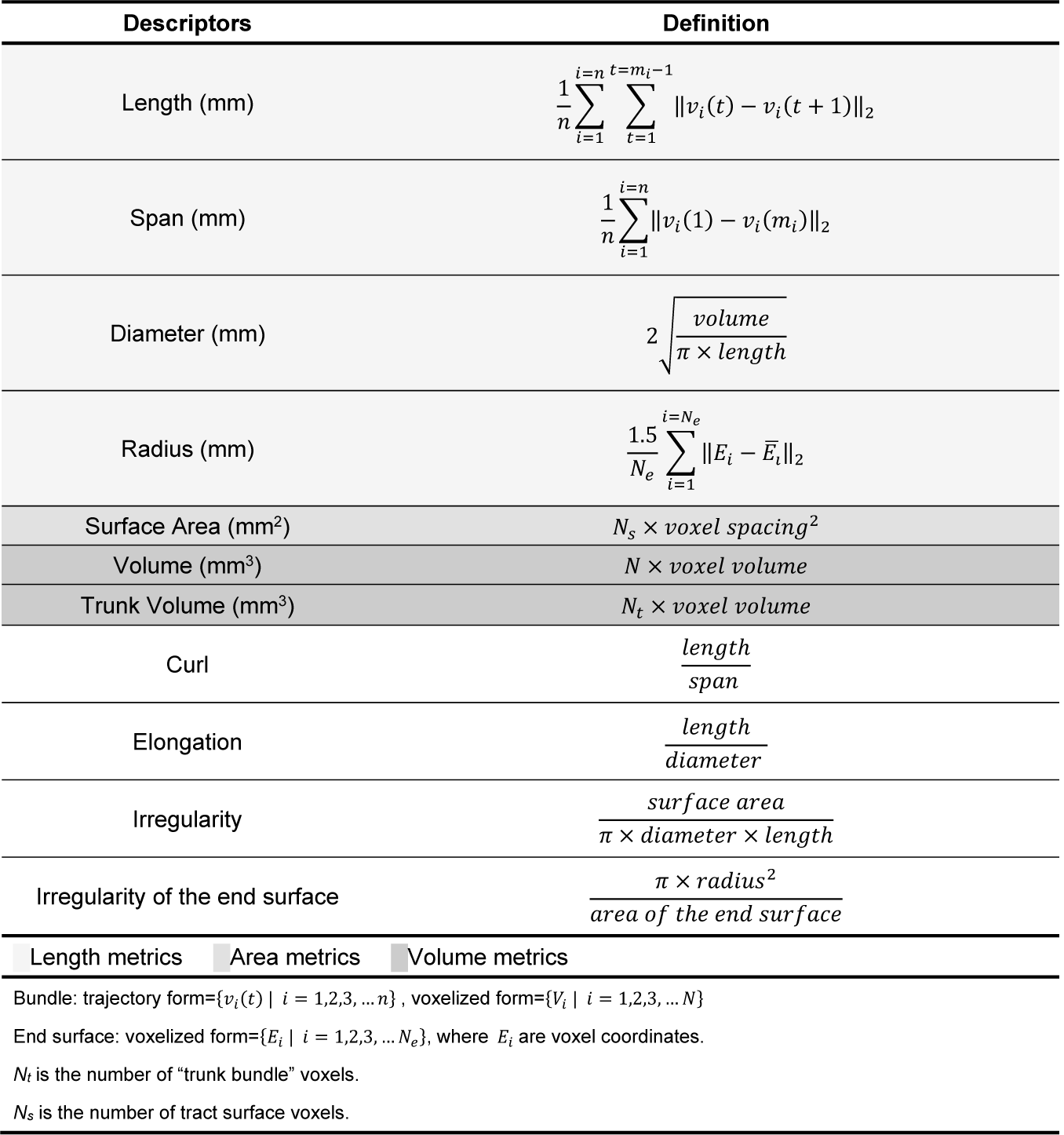
List of shape descriptors and their definition

We mapped 14 association pathways, including the left and right arcuate fasciculus (AF), cingulum (C). frontal aslant tract (FAT), inferior fronto-occipital fasciculus (IFOF), inferior longitudinal fasciculus (ILF), superior longitudinal fasciculus (SLF), and uncinate fasciculus (UF). The SLF here included “SLF II” and “SLF III,” as they often form a continuous sheet structure together. “SLF I” was not included because it is often separated from the other two SLF bundles and closely sided with cingulum. The starting region of the fiber tracking (a.k.a. the seeding region) was defined using the corresponding white matter regions in the HCP842 tractography atlas (Yeh et al., 2018) (nonlinearly registered to the subject’s native space). To cope with the reproducibility problem in tractography, we saturated the tracking parameters using a random generator to select a combination of fiber tracking parameters within a working range. The tracking parameters included the anisotropy threshold, angular threshold, step size (a.k.a. the propagation distance). The anisotropy threshold was randomly selected between 0.5 and 0.7 of the Otsu’s threshold (Otsu, 1979). The angular threshold was randomly selected between 15 to 90 degrees. The step size was randomly selected between 0.5 to 1.5 voxel distance. The random generator was based on a uniform distribution to select a value from the above parameter range. For each of the 14 association pathways, we initiated 5,000,000 tracking iterations, with each iteration having a unique sample of the parameter combination. The fiber tracking was conducted using a deterministic fiber tracking algorithm (Yeh et al., 2013).

### Automatic track recognition

The generated 5,000,000 tracks were further filtered by automatic track recognition. The recognition was based on the shortest Hausdorff distance with the trajectories of the HCP842 tractography atlas (Yeh et al., 2018). For each track, we calculated its Hausdorff distance with each trajectory in the atlas (nonlinearly wrapped to the subject space), and the shortest distance will correspond to an atlas trajectory that allowed us to identify the anatomical nomenclature of the track. After recognition, we used 16 mm as the maximum allowed threshold for the shortest distance. This threshold will help to remove tracks with substantial deviation from the known atlas trajectories. In this study, the recognition did not find the right arcuate fasciculus in two subjects (both test and retest scans). Further investigations into these two subjects found that the initial fiber tracking did generate numerous pathways from the right arcuate fasciculus area, but subsequent tract recognition categorized them as right superior longitudinal fasciculus because the trajectories did not reach the right superior temporal lobe. The analysis of the right arcuate fasciculus in this study thus excluded these two subjects.

### Topology-informed pruning

All recognized trajectories were then summed up and pruned by topology-informed pruning (Yeh et al., 2019). A total of 20 pruning iterations was conducted. The pruning method calculated the voxel-wise streamline density and used low-density voxels to identify noisy tracks. Since the white matter pathway tends to form a bundle or a sheet, removing noisy tracks could increase the accuracy of the overall tracking results. We pruned all association pathways except for the right arcuate fasciculus. In one subject, the right arcuate fasciculus was too thin, and the pruning iteration was reduced to 10 to avoid pruning out all tracks. After pruning, the shape characteristics were quantified using the following shape analysis.

### Shape analysis

In computer vision, shape analysis quantifies 2D or 3D objects using shape descriptors such as curl, elongation, roundness for 2D or 3D shapes (Russ, 2002)(Page 513). Some of them can be directly translated to tractography, whereas others had to be modified to consider that tractography is a set of trajectories in the 3D space. The following section details how each descriptor was calculated from tractography streamlines:

A fiber bundle is a set of streamline trajectories that can be represented as 3D coordinate sequences: {*v*_*i*_(*t*) | ***i*** = 1,2,3, … *n*}. Here *n* is the total number of tracks, *v*_*i*_(*t*) is a sequence of 3D coordinates representing the trajectory of a track. *t* is a discrete variable from 1 to *m*_*i*_, where *m*_*i*_ is the number of the coordinates. The length of a fiber bundle is thus defined as follows:

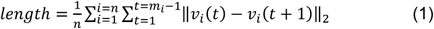

The span is defined as:

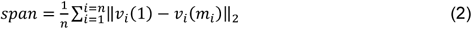

Following the conventional definition (Russ, 2002), curl is defined as:

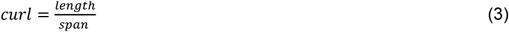

Curl has a range of [1, ∞). A track bundle with a big curl value tends to have a curvy shape, whereas a straight line has a curl value of 1.

Then we voxelized tracks to carry out further shape analysis. All trajectories were first resampled so that for any two consecutive coordinates in any track, their distance was smaller than the voxel size. This resampling allowed us to directly “voxelize” tracks by rounding up all coordinates and removing repeated coordinates. To minimize discretization error, we multiplied track coordinates by two before rounding up, and any further metrics calculation will consider this scaling effect. The voxelized tracks could be represented by a set of unique voxel coordinates denoted as *T* = {*V*_*i*_ | *i* = 1,2,3, … *N*}, where *N* is the total number of unique voxel coordinates. The total track volume could be estimated by the following:

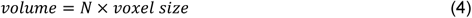

Note that due to our previous scaling, the voxel size was 4^3^ times smaller than the raw DWI voxel size. The bundle diameter was then approximated using a cylinder model:

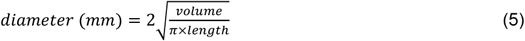

The diameter can be used to calculate elongation as a shape metric:

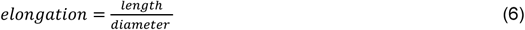

This definition follows the one used in computer vision. To calculate track surface area, we converted the track voxel set *T* to a 3D volume *V*(*x, y, z*), whereas *V*(*x, y, z*) = 1 if *V*(*x, y, z*) ∈ *T* and 0 otherwise. This 3D volume enabled us to use morphology operation to identify the “surface voxels,” which had non-zero values and connected to at least one zero-valued neighboring voxel. The surface area was then estimated as follows:

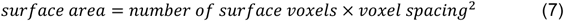

It is noteworthy that the surface area here will include the innervation area. Based on a cylinder model, we propose a new descriptor called “irregularity,” which is defined as

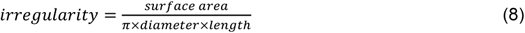

Irregularity is conceptually similar to convexity and concavity. It is the opposite of compactness or roundness defined in computer vision (Russ, 2002). A surface area much larger than the expected cylinder surface suggests higher shape irregularity.

The rest of the shape analysis then utilized the two end surfaces of a track bundle. The end surfaces were determined by anisotropy and angular threshold used in the fiber tracking. One obstacle for end surface analysis was that the coordinates of a track could be sequenced in two opposite directions (antegrade or retrograde), and correctly grouping the endpoints into two “end surfaces” required additional clustering steps. To this end, we used k-means clustering algorithm with *k*=2 and modified it to ensure that the two endpoints of the same track were placed in the different clusters. The clustering started with all *v*_*i*_(1) assigned to cluster 1 and all *v*_*i*_(*m*_*i*_) assigned to cluster 2. Then the mean coordinate for each cluster was computed, and we clustered the endpoints of each track again using their distance to the mean coordinates. The above steps were repeated until there was no cluster change for all the endpoints. This resulted in two sets of coordinates (one for each end surface). The coordinates of the clustered endpoints were then rounded up to remove repeat voxel coordinates. This generated two unique sets of discrete voxel coordinates: *E*_1_ and *E*_2_, each of them denoting the voxelized end surfaces of the track bundle. We further checked the mean coordinates of *E*_1_ and *E*_2_ and figure out which of the x-, y-, or z-dimension has the largest distance between the mean coordinates. Without loss of generality, we assigned *E*_1_ to be the end surface that had a larger coordinate value in at this dimension (posterior or superior end of a bundle). The area of *E*_1_ or *E*_2_ was then calculated as follows:

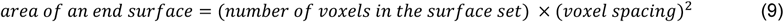

The area was calculated separately for each of the end surfaces. To quantify the extent of an end surface, we define a new descriptor called the radius, which was calculated by modeling the end surface as a circle. Assuming we have a uniform endpoint distribution in a disk, the mean distance to the center will be 2/3 the radius. Thus the estimated radius is 1.5 of the mean distance:

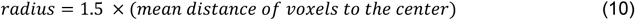

We also introduce irregularity for the end surface. The irregularity was calculated as follows:

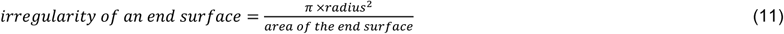

The irregularity of a circle will be 1, whereas any protrusion or intrusion will increase the irregularity. Last, the end surface coordinates were used to define the “trunk” of a bundle. We first converted *E*_1_ and *E*_2_ into two 3D volumes of 0-1 valued voxels, respectively. The converted volumes were then analyzed by 3D connected component analysis to isolate the largest region of the surface. The two generated regions were then used as two regions of interest to isolate the main trunk of the fiber bundle and calculate its volume.

For each shape descriptor, the test-retest reliability was calculated using one-way random, single measures intraclass correlation (ICC 1-1). The median value of descriptors from 14 bundles was calculated as an overall indicator of the performance. The between-subject variations of each descriptor were quantified using the absolute deviation from the median further divided by the median to facilitate comparison.

### Computation resources

The source code is available at http://dsi-studio.labsolver.org with documentation to ensure the reproducibility of this study. The analysis was conducted on the “Bridges” supercomputer at Pittsburgh Supercomputing Center provided through the XSEDE resource (Towns et al., 2014). The Bridges supercomputer network has 752 computation nodes, and each node has two 14-core Intel Haswell CPU at 3.3 GHz. XSEDE is an NSF-funded (US) program providing supercomputer resources shared by multiple research groups. This study used start-up allocation to accomplish the computation task, and DSI Studio package was available through its “singularity” container (Kurtzer et al., 2017) by pulling the docker container at docker://dsistudio/dsistudio. The batch processing was realized using the command line interface (http://dsi-studio.labsolver.org/Manual/command-line-for-dsi-studio). The computation time for each pathway in one subject was around 3∼5 minutes. The data transfer time and the waiting time at job queues were substantially longer (hours to a day). The total size of the tractography generated (44 subjects × 14 pathways) was 168 GB (in trk.gz format).

## Results

### Augmented fiber tracking

Figure 2a shows the tracking result of the first subject, including the arcuate fasciculus (AF), cingulum (C). frontal aslant tract (FAT), inferior fronto-occipital fasciculus (IFOF), inferior longitudinal fasciculus (ILF), superior longitudinal fasciculus (SLF), and uncinate fasciculus (UF) presented in the left, right, anterior, and superior views. Only the association pathways in the left hemisphere are shown here to facilitate comparison. The tractography matches the known anatomical trajectories of the human association pathways, suggesting the feasibility of the automatic track recognition to obtain clean results without manual intervention. Figure 2b further shows the left arcuate fasciculus of all 44 subjects, including their test-retest results generated from automatic track recognition. The tractography of the repeat scan is placed immediately on the right of the first scan. The tracking results show C-shaped bundles that match the anatomy of the arcuate fasciculus. This qualitative evaluation suggests that the augmented fiber tracking could provide grossly consistent tractography results in test-retest scans.

**Figure 2.**
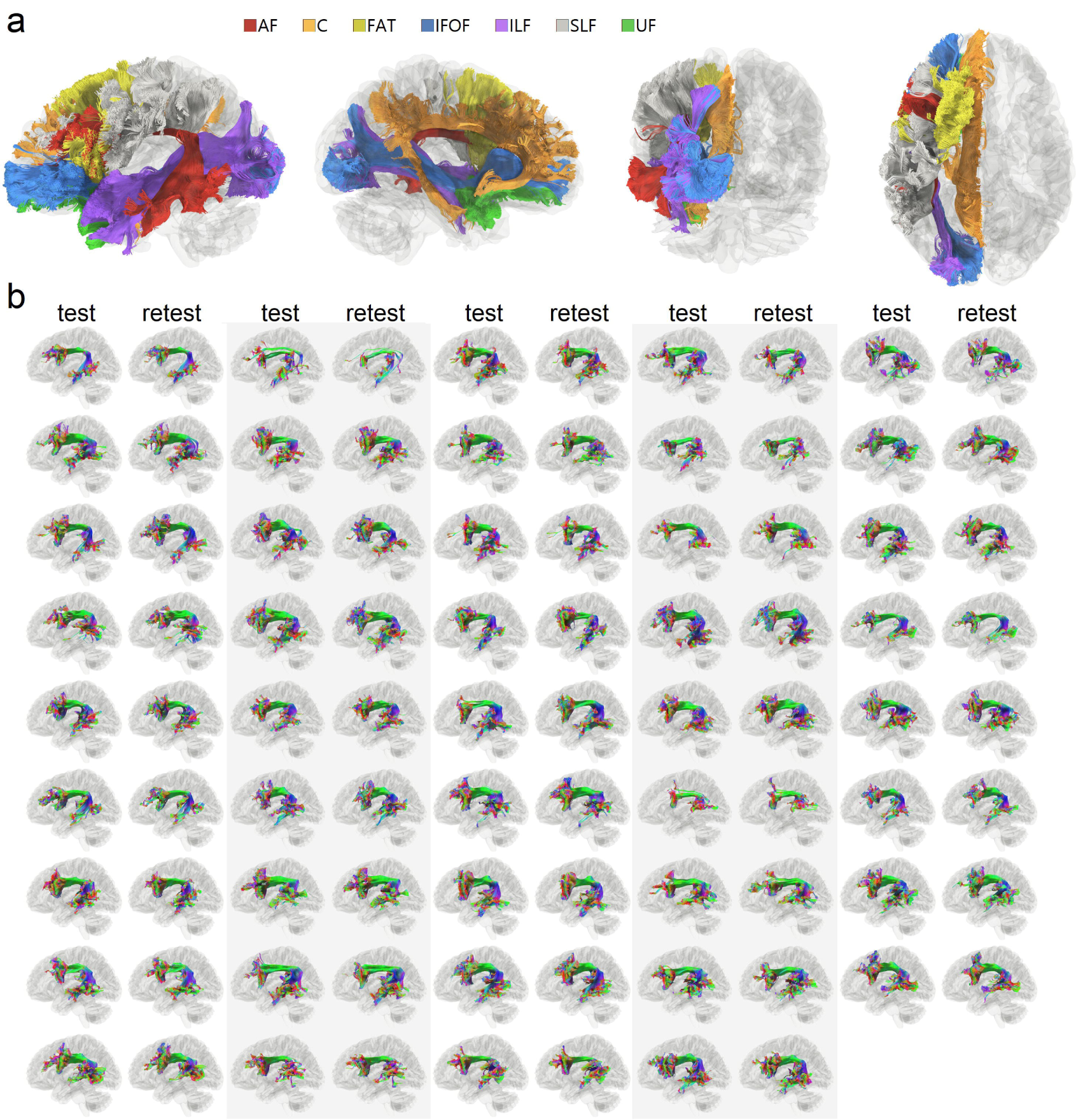
Qualitative evaluation of association pathways generated using augmented fiber tracking. (a) Seven association pathways in the left hemisphere of a subject are automatically tracked. The tracking results seem to match well with the known neuroanatomical structures of the association pathways. (b) The result of arcuate fasciculus tractography of all test-retest scans (n=44×2) mapped using augmented fiber tracking. All results show similar C-shaped bundles. The test-retest results are grossly consistent, suggesting the reliability of the method for high throughput analysis.

Figures 3a and 3b further present the test-retest results of the arcuate fasciculus tractography in more detail. We selected three best (Fig. 3a) and three worst (Fig. 3b) performers from our subject pool, as quantified by the differences in the volume between the test-retest scans. The tractography in Figure 3a shows high consistency in the fiber trajectories. The topological pattern of the core bundle is almost identical, though minor differences can still be observed at the details. Figure 3b shows tractography from the three worst performers in the test-retest scans. Although at their worst, the overall tractography still also presents decent consistency. Most of the differences are located in the branches, whereas the core trajectories are still highly consistent.

**Figure 3.**
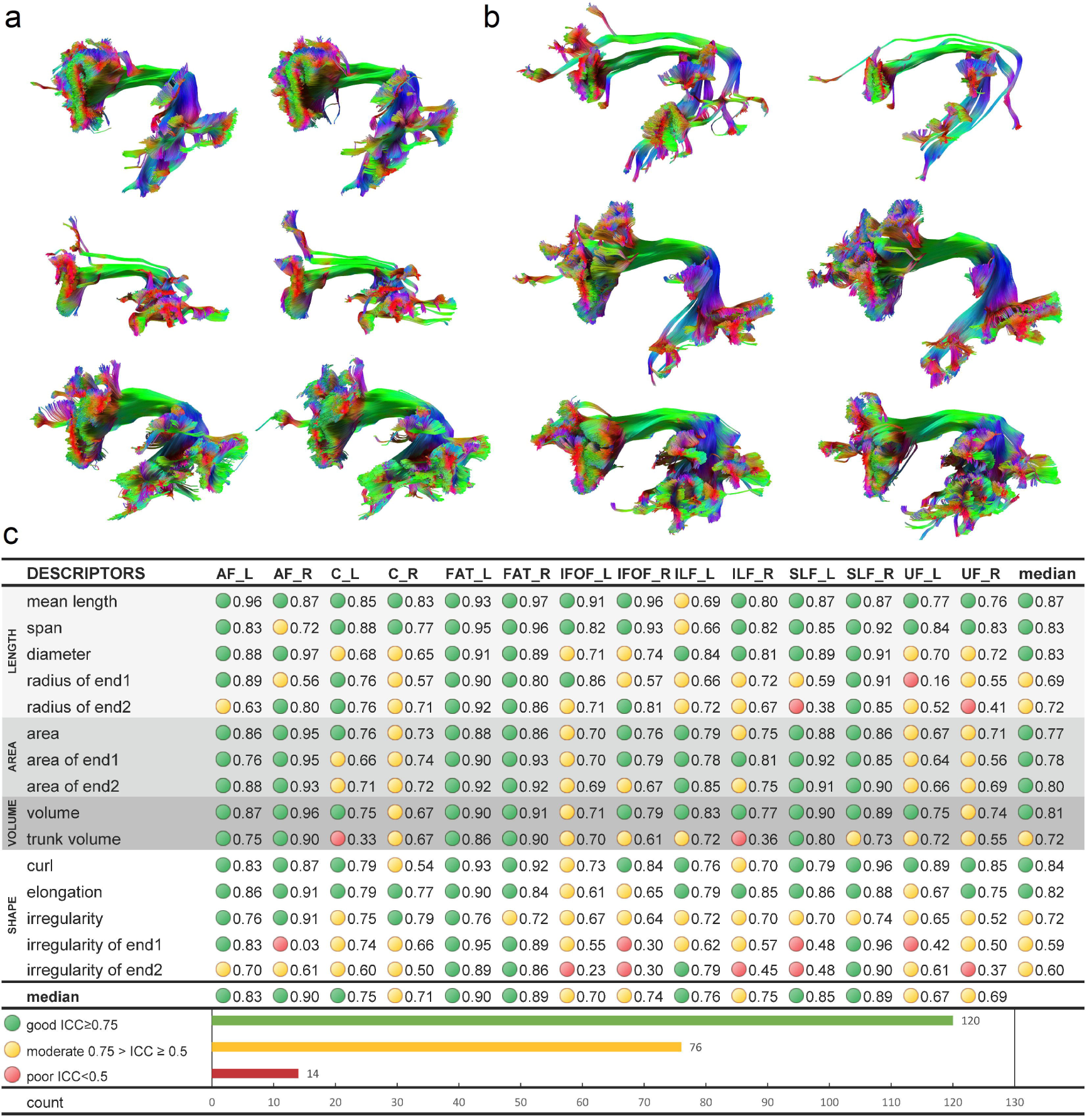
Reproducibility of tractography and reliability of the shape descriptors. (a) Three subjects with the best performing test-retest results are selected for their small differences in test-retest volume. The tractography from test and retest scans is of high similarity, while the unique structural characteristics of each subject are preserved. (b) Three subjects with the worst performing test-retest results are selected as a comparison. Even at its worst, the fiber tracking still achieves decent consistency between test-retest scans and preserves the structural characteristics of each subject. (C) The test-retest reliability of the shape descriptors is quantified by intraclass correlation (ICC). The majority of the shape descriptors show moderate (>0.5) to good (>0.75) reliability. The median ICC values for all descriptors are greater than 0.5, while poor reliability (<0.5) still presents in around 6% of the application scenarios.

### Test-retest reliability

Figure 3c lists the intraclass correlation (ICC) of shape descriptors for each bundle. The shape descriptors can be categorized into length metrics (light gray), area metrics (gray), volume metrics (dark gray), and shape metrics (white). Good reliability (ICC≥0.75) is labeled by a green circle, and moderate reliability (0.75>ICC≥0.5) is labeled by a yellow circle. Poor reliability (ICC<0.5) is marked by red. Out of 210 bundle-descriptor entries, 120 of them (57.1%) have good reliability, 76 of them (36.2%) have moderate reliability, and 14 of them (6.7%) have poor reliability. More than 90% of the scenarios have moderate to good reliability, suggesting overall good reliability of the shape descriptors. All descriptors have a median ICC value greater than 0.5, and the length metrics perform the best, with a median value of ICC around 0.8. The area and volume metrics are the next, showing the median values of ICC around 0.7∼0.8. The shape metrics moderate to good reliability, with curl and elongation performing the best, and irregularity the last. There are poor reliability scenarios in radius, trunk volume, and irregularity that requires precautions. These metrics can have outstanding reliability (ICC>0.9) for some bundles and poor reliability (ICC < 0.5) in the others. This indicates that the application of these three shape descriptors still requires additional precautions to avoid poor reliability conditions.

### Normative distribution of shape descriptors and left-right asymmetry

Figure 4a shows representative examples of large and small metrics values using the left arcuate fasciculus selected from the subject pool. The mean and standard deviation values of the shape descriptors are listed and color-coded in Fig. 4b. Only right-handed subjects were included in all the following analyses. In Fig. 4b, the red color represents a relatively higher value compared with other association pathways. For example, the length of the inferior fronto-occipital fasciculus (IFOF) is marked by red, suggesting their longest length among all association pathways. Similarly, the frontal aslant tract (FAT) has the largest diameters, and the left cingulum (C) has the largest surface area. The left superior longitudinal fasciculus (SLF) has the highest topological irregularity.

**Figure 4.**
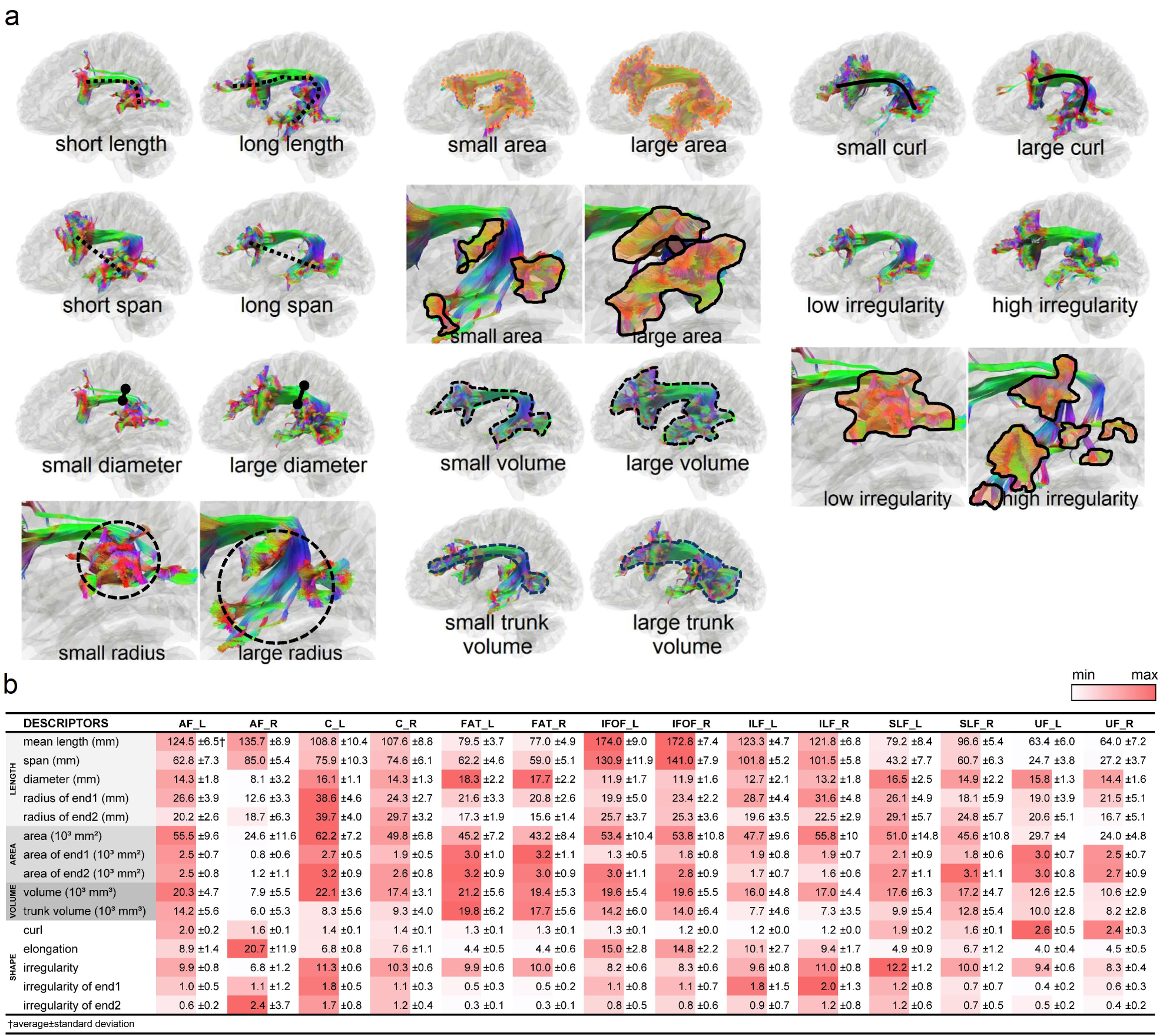
(a) Representative cases of shape descriptors are shown using the left arcuate fasciculus as an example. (b) Mean values and standard deviations of shape descriptors are listed for each association pathway. The red colors are those with relatively large values in comparison with other pathways.

We further plot the distributions of length metrics in Fig.5 for each of the association pathways. The end surface 1 and 2 for each bundle are labeled on the tractogram to the top. In each of the box plot figures, the two circles on the right upper corner represent the test-retest reliability of the measures, as listed in Fig.3c. Green color indicates good test-retest reliability (ICC≥0.75), yellowish color indicates moderate reliability (0.75>ICC≥0.5), and red color indicates poor reliability (ICC<0.5). The distributions for the left side bundle are colored by blue, whereas the right colored by red. Paired t-tests were used to test the left-right differences (null hypothesis: the left-right differences are zero). The p-value results are presented with significance marks (*< 0.05, **<0.01, ***< 0.001), and the percentage differences are also calculated by 100%×(*a*-*b*)/*a*, where *a* is the quantity of the dominance side.

The largest and most significant left dominance can be found in the arcuate fasciculus (AF) in the diameter and radius of its posterior end surface at the temporal lobe. On the next, superior longitudinal fasciculus (SLF), cingulum (C), and uncinate fasciculus (UF) show moderately left dominance at 10∼20% in diameter. Superior longitudinal fasciculus (SLF) and cingulum (C) further shows significant left dominance in the radius of their innervation surfaces. The left frontal aslant tract (FAT) shows a slightly larger radius of the innervation region at the inferior frontal lobe. In comparison, the inferior fronto-occipital fasciculus (IFOF) and inferior longitudinal fasciculus (ILF) shows right-dominance only in the radius of the end surfaces with no significant difference in the diameter.

Figure 6 further shows the distributions of area and volume metrics for the association pathway bundles. The arcuate fasciculus (AF) shows a large left-dominance in area and volume metrics greater than 50%. On the next, cingulum (C), and uncinate fasciculus (UF) show moderately left dominance at ∼20% in area and volume. The frontal aslant tract (FAT) shows only a slightly larger volume in the left hemisphere (14.8%). In comparison, inferior longitudinal fasciculus (ILF) shows moderate right-dominance in the area with no significant difference in the volume. The inferior fronto-occipital fasciculus (IFOF) shows right-dominance in the area of the posterior end surface. The superior longitudinal fasciculus (SLF) has a more complicated lateralization profile, with left dominance in tract area and right dominance at the anterior innervation region and trunk volume. Findings from Figs. 5 and 6 show an overall trend of left-dominance in the arcuate fasciculus (AF), cingulum (C), frontal aslant tract (FAT), and uncinate fasciculus (UF), and right dominance in the inferior fronto-occipital fasciculus (IFOF) and inferior longitudinal fasciculus (ILF). The superior longitudinal fasciculus (SLF) has mixed lateralization with different metrics showing either left or right dominance.

**Figure 5.**
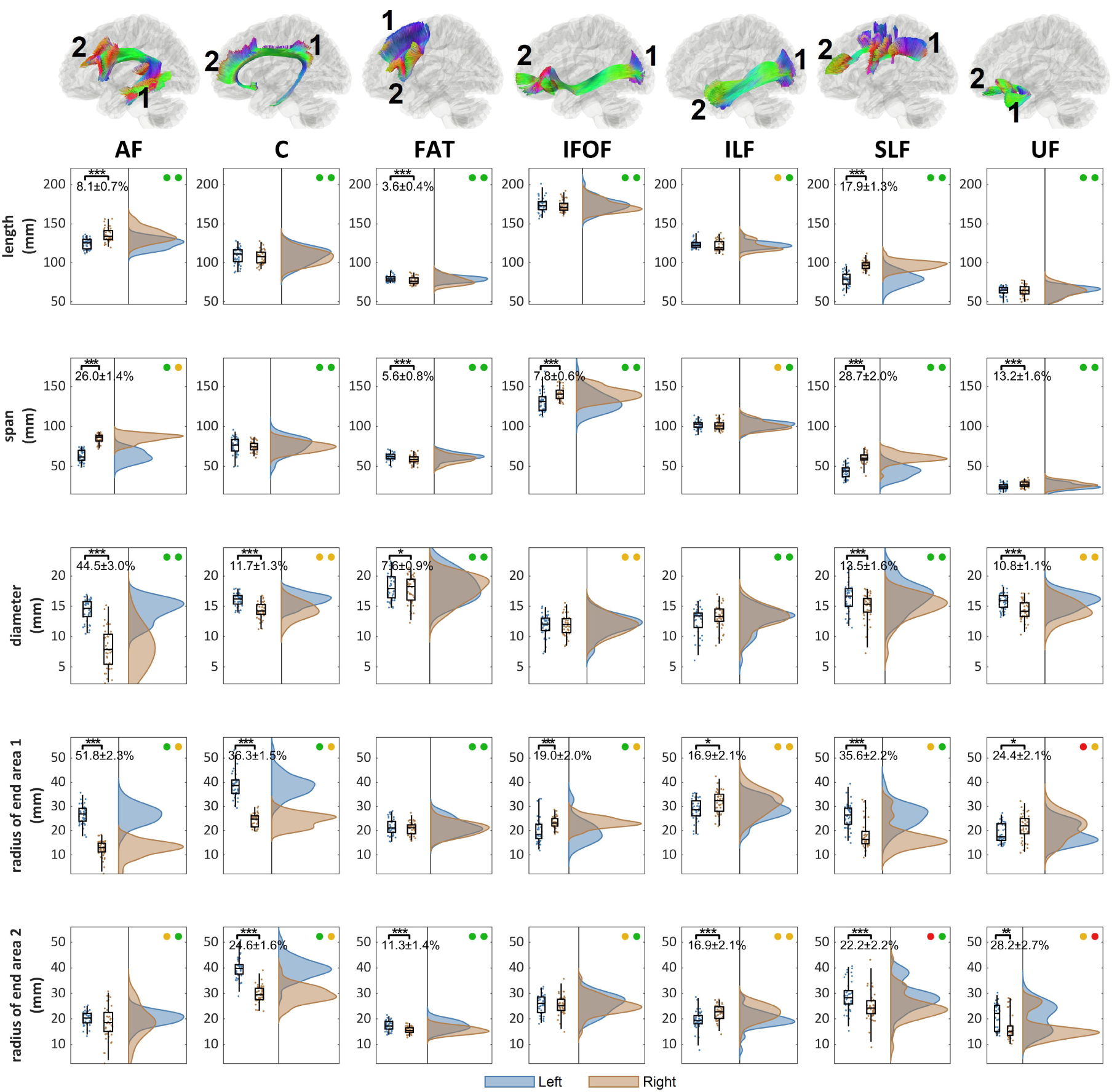
The distribution of the length metrics and their left-right differences in the association pathways. The location of the end surface 1 and 2 are annotated for each bundle. The association pathways present different significance level of the left-right differences (p-value: *** < 0.001, **< 0.01, *< 0.05). The test-retest reliability of the metrics for the left and right bundle is presented by colored circles (green: ICC≥0.75, yellow: 0.75>ICC≥0.5, red: ICC<0.5). AF, C, FAT, SLF, and UF present an overall left dominance in either the diameter or radius, whereas IFOF and ILF present right dominance.

**Figure 6.**
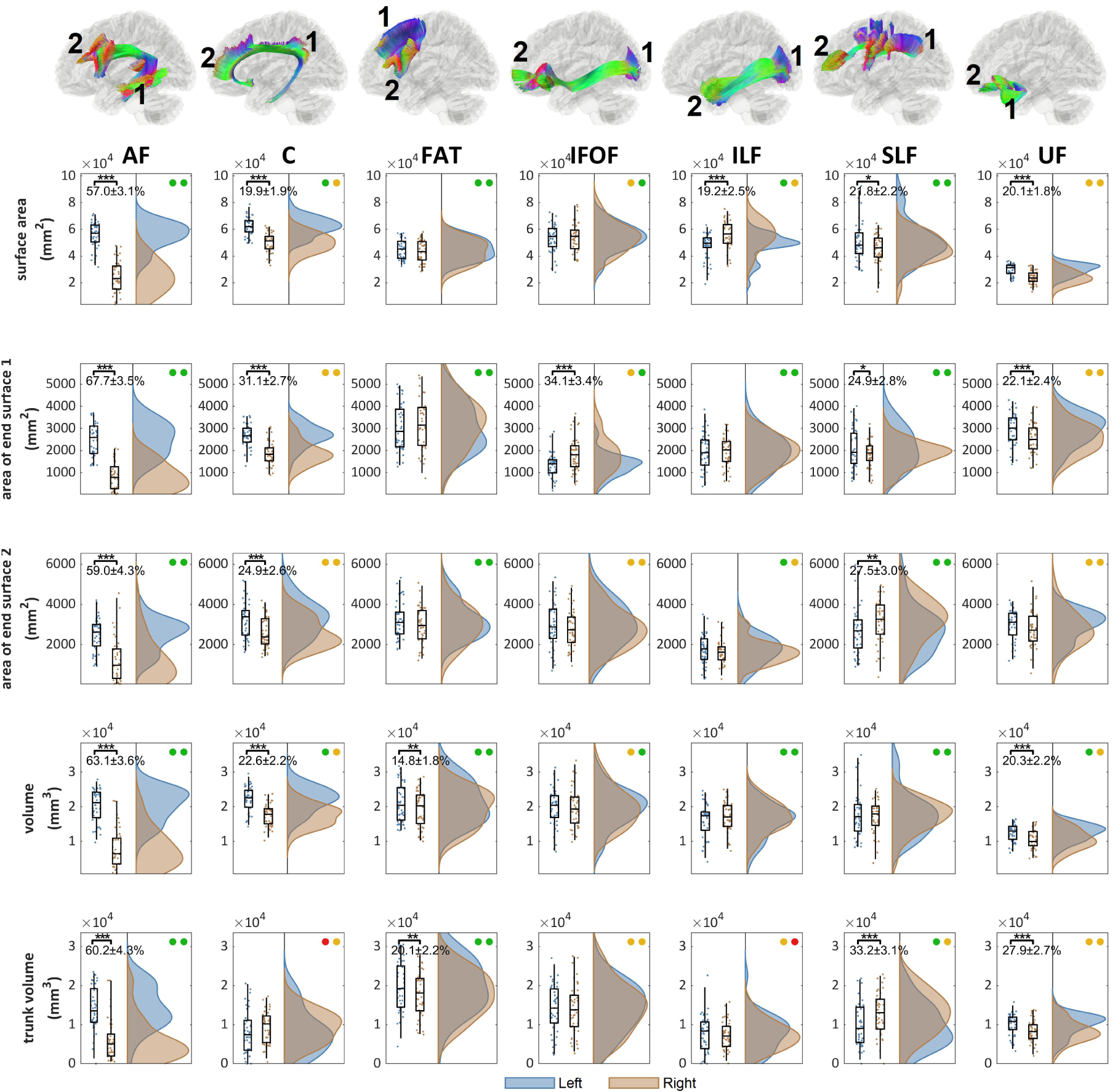
The distributions of the area and volume metrics and their left-right differences in the association pathways. The location of the end surface 1 and 2 are annotated for each bundle. The left-right differences are tested (p-value: *** < 0.001, ** < 0.01, * < 0.05). The test-retest reliability of the metrics for the left and right bundle is presented by colored circles (green: ICC≥0.75, yellow: 0.75>ICC≥0.5, red: ICC<0.5). AF, C, FAT, and UF shows significant left dominance in either area or volume metrics, whereas IFOF and ILF show significant right dominance. SLF presents a mixed lateralization profile with either right or left dominance in different metrics.

Figure 7 shows the distribution of shape metrics for the association pathways. The differences between left and right distribution are quantified using Cohen’s d. While all pathways present significant left-right differences in different shape metrics, the irregularity metric presents the most significant and largest left-right asymmetry. The arcuate fasciculus (AF), cingulum (C), superior longitudinal fasciculus (SLF), and uncinate fasciculus (UF) shows substantial left dominance in the irregularity (paired t-test p-value < 0.001, d > 1.5), while inferior longitudinal fasciculus (ILF) shows right dominance (paired t-test p-value < 0.001, d=1.76).

**Figure 7.**
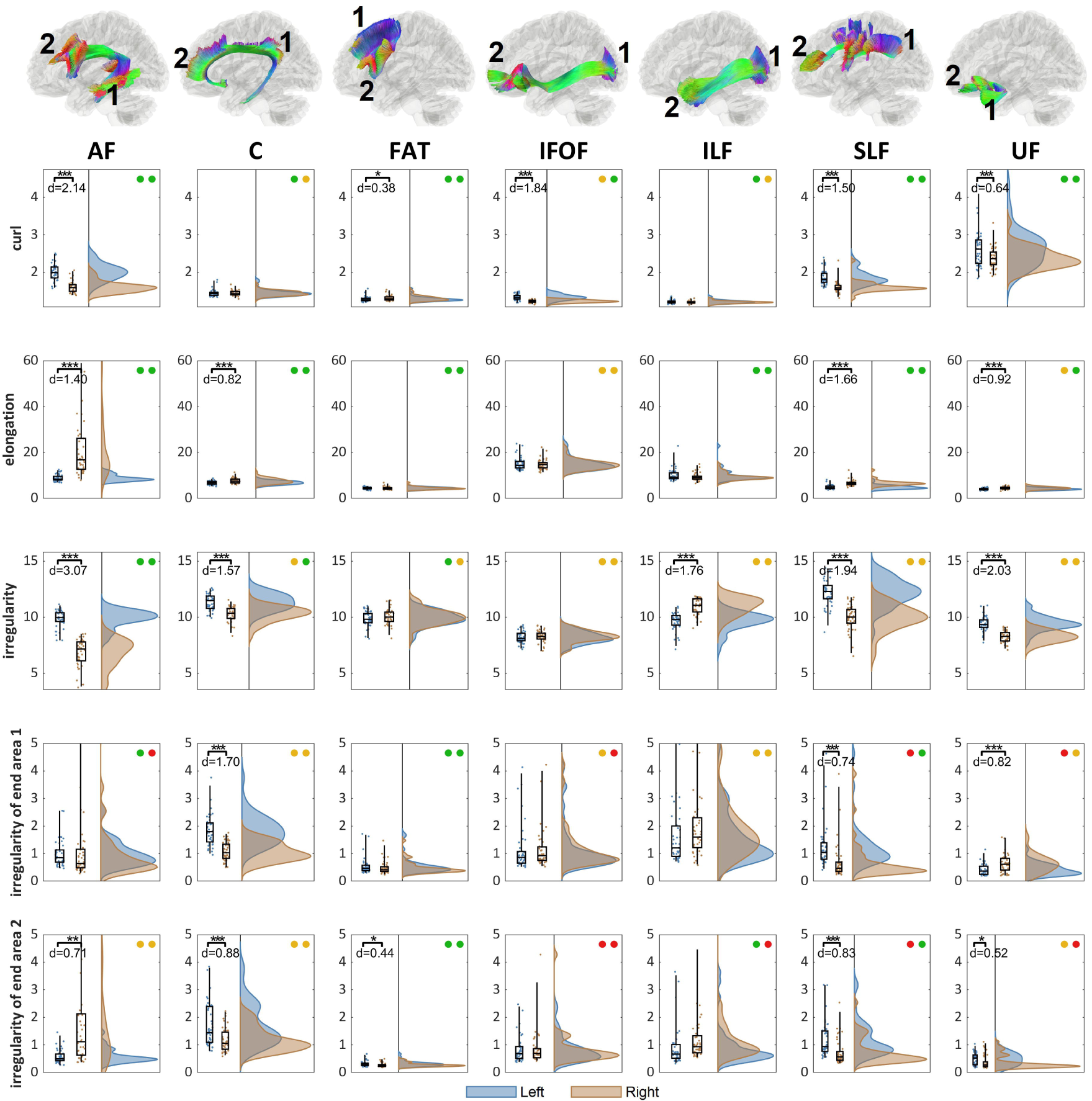
The distributions of the shape metrics and their left-right differences in the association pathways. The location of the end surface 1 and 2 are annotated for each bundle. The left-right differences are tested (p-value: *** < 0.001, ** < 0.01, * < 0.05) and effect size (Cohen’s d) with test-retest reliability presented as colored circles (green: ICC≥0.75, yellow: 0.75>ICC≥0.5, red: ICC<0.5). All pathways present significant lateralization at different shape metrics. The overall irregularity shows the large left-dominance at AF, C, SLF, UF, and right dominance at ILF, suggesting their prominent left-right differences in bundle topology.

### Between-subject variations

Figure 8 shows the between-subject variations using the absolute deviation. The absolute deviation was calculated by the absolute difference from the median to evaluate the dispersion of the shape descriptors between subjects. The deviation was further divided by the median value of the bundle to facilitate comparison. Furthermore, the overall median value of all bundles is plotted by a blue vertical line, whereas the first and third quantiles are plotted by a red line. As shown in Fig. 8, the length, span, and diameters have small between-subject differences, mostly less than 10% deviations. The variations in diameter are larger for the right arcuate fasciculus (AF_R), likely due to its smallest diameter among all association pathways. The radius and surface also have a similar variation level, with the majority of the deviations lower than 20%. A much larger between-subject variation can be observed for the are of the end surfaces, mostly ranged between 10∼40% in the absolute deviation. The overall results suggest that the “layout” of the association pathways seems not to vary a lot across subjects, as shown by low between-subject variation in length, span, diameter, and radius. In contrast, the innervation region has a considerable between-subject variation that may account for most of the individual differences in white matter structure.

**Figure 8.**
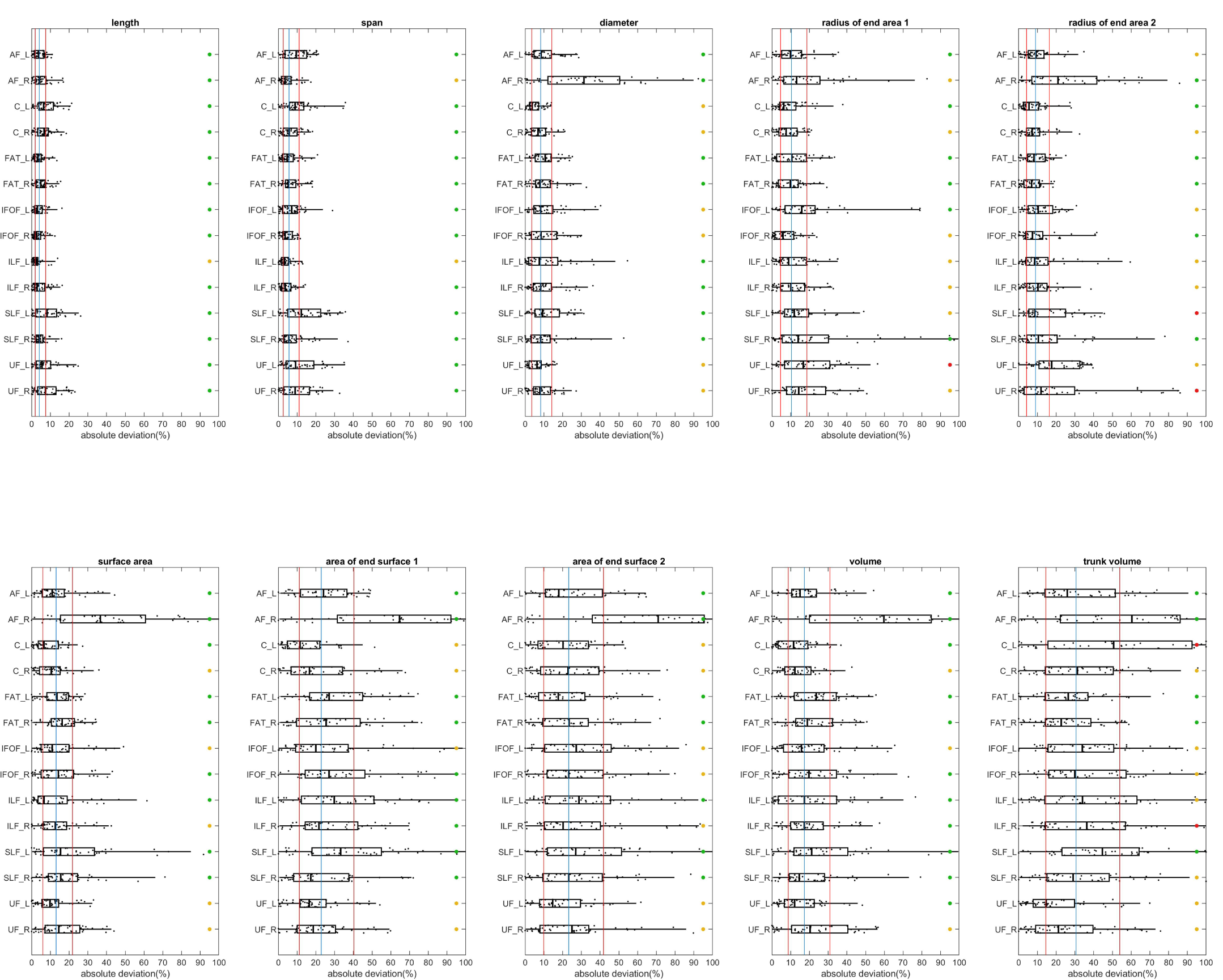
Between-subject variations of the length, area, and volume metrics in the association pathways. The variations are quantified by absolute deviation. The blue vertical line marks the median of deviation values of all bundles, whereas the two red vertical line marks the first and third quantiles. The test-retest reliability is labeled by colored circles (green: ICC≥0.75, yellow: 0.75>ICC≥0.5, red: ICC<0.5). All length metrics have relatively smaller between-subject variation, and the area and volume metrics show a slightly larger between-subject variation. The end surfaces show a greater deviation of more than 20%. The overall results suggest that the “layout” of the association pathways seems not to vary a lot across subjects, while the innervation region has a considerable between-subject variation.

## Discussion

Here we conducted shape analysis on human association pathways and confirmed its reliability in a test-retest dataset. We derived the distribution of shape descriptors to elucidate lateralization and between-subject variations. The results revealed an overall left dominance in arcuate fasciculus, cingulum, uncinate, and frontal aslant tract, with the largest lateralization found in the arcuate fasciculus. Cingulum and uncinate fasciculus showed moderate lateralization in either diameter, area, or volume, while the frontal aslant tract showed small lateralization. Right dominance was found in inferior fronto-occipital fasciculus and inferior longitudinal fasciculus. Although there was a widespread left-right asymmetry in all association pathways, the detail lateralization profile varied substantially across bundles, and not all bundles share the same lateralization pattern.

The lateralization found in this study is not new to the neuroscience field. For example, studies have shown lateralization in the arcuate fasciculus (Lebel and Beaulieu, 2009; Vernooij et al., 2007) and the inferior longitudinal fasciculus (Panesar et al., 2018), yet our findings revealed a more sophisticated profile in lateralization. A bundle could have left dominance in one metric and right dominance in another, and a comprehensive profile covering all metrics is needed to investigate the asymmetry thoroughly.

In addition to lateralization, the between-subject variation quantified in this study gave us a glimpse into how white matter structures variate across the population. Our analysis showed that the between-subjects variation was small in length metrics such as length, span, diameter, and radius, whereas the area of the end surfaces had a much larger variation. While the length and span did not vary much (< 10% deviation), the area of the innervation region had a median deviation of 24%, implying a considerable variation in how white matter bundle innervates at the cortical surface.

### Technical challenges and limitations

This study still has limitations. Good test-retest reliability in shape analysis only implies the robustness of the algorithm. It does not necessarily guarantee that the results are always correct. The fiber tracking algorithm still has the issue of false-positive and false-negative results. For deterministic fiber tracking, false-negative results are more common, as the ability to capture more delicate branches depends on the spatial resolution and the sensitivity of the data acquisition. There are possibilities that a minor branch was left undetected in both test and retest scans due to the limitation of acquisitions. Another challenge is the accuracy of automatic track recognition. Several diffusion MRI tools are providing similar functionality using atlases, cortical regions, or bundle clustering (Garyfallidis et al., 2018; Warrington et al., 2020; Wasserthal et al., 2018; Watanabe et al., 2018; Yeatman et al., 2012). The track recognition in this study used a tractography atlas as the only reference and did not use cortical regions as the inclusion or exclusion criteria. The purpose was to avoid issues related to cortical parcellations. Whether this approach provides better or worse performance requires further investigation. The shape descriptors quantified in this study can be further improved. The volume and area metrics calculated in this study were calculated using voxelization and thus subject to discretization error. A better quantification strategy is to use surface triangularization to achieve better accuracy in volume calculation. For the end surface of the bundle, the tract termination can be projected to the gyral surface to achieve a more robust quantification. Last, we only have 44 subjects included in the analysis because of the long computation time needed for automatic fiber tracking. To further investigate between-subject differences, we are planning a future population-based study to include all 1065 HCP subjects and describe the normative variation of white matter structures.

Nonetheless, there are encouraging reproducibility achieved in this study. We showed that a combination of parameter saturation, automatic track recognition, and topology-informed pruning could provide good reproducibility. The derived metrics further achieved moderate to good test-retest reliability. By integrating with shape analysis, diffusion MRI has a new option for white matter analysis. It can be used in neurological, psychological, and psychiatric studies to investigate the correlation between white matter architecture correlates and abnormal brain functions, with a hope to decipher how structure defines brain functions.

## Acknowledgments

The research reported in this publication was partly supported by NIMH of the National Institutes of Health under award number R56MH113634. The content is solely the responsibility of the authors and does not necessarily represent the official views of the National Institutes of Health. This work used the Extreme Science and Engineering Discovery Environment (XSEDE), which is supported by the National Science Foundation grant number ACI-1548562. Data were provided by the Human Connectome Project, WU-Minn Consortium (Principal Investigators: David Van Essen and Kamil Ugurbil; 1U54MH091657) funded by the 16 NIH Institutes and Centers that support the NIH Blueprint for Neuroscience Research; and by the McDonnell Center for Systems Neuroscience at Washington University.

